# Thirty-five experimental fisheries reveal the mechanisms of selection

**DOI:** 10.1101/141259

**Authors:** Sébastien Nusslé, Andrew P. Hendry, Roland A. Knapp, Michael T. Bogan, Anna M. Sturrock, Stephanie M. Carlson

## Abstract

Fisheries have been described as large-scale evolutionary experiments; yet such “experiments” tend to be poorly replicated and therefore lack the predictive power essential for designing appropriate management strategies to minimize the effects of fisheries-induced selection. Large-scale removal of non-native trout from 35 montane lakes in California provided repeated experimental fisheries that allowed us to explore how environmental parameters affect the three potential contributors to overall selection: the fitness-trait correlation, trait variability, and fitness variability. Our results demonstrate that fishing rapidly altered the size structure of harvested populations, and that the magnitude of change was primarily driven by the fitness-trait correlation (net selectivity). Fishing-induced selection was repeatable overall but was also influenced by environmental (lake size and quality) and demographic (size structure) parameters. Decomposing fishing-induced selection into its key components can improve the management of stocks experiencing fishing-induced selection by identifying the drivers of selection and therefore the appropriate target for management.

## Text

By targeting the oldest, largest, and most fecund individuals, size-selective fishing can reduce recruitment and yield ^1,2^ and induce evolutionary changes that negatively influence stock productivity, resiliency, and recovery ^3,4^. However, two major issues have hindered efforts to incorporate fishing-induced selection into conservation plans and management strategies ^5–7^. First, studies of fishing-induced selection are typically not well replicated within a species, and therefore lack the potential to test for causal drivers through comparative analyses ^8^. Second, selection can be shaped by three key components (the fitness-trait correlation, trait variability, and fitness variability) ^9,10^ that have not been considered individually in fisheries science, and yet have different implications for management (see below). We circumvented these previous limitations through a study of fishing-induced selection in 35 independent populations of fishes (brook trout, *Salvelinus fontinalis*; and rainbow trout, *Oncorhynchus mykiss*).

The study populations were fished to extirpation through a large-scale non-native fish removal as part of a habitat restoration and endangered species recovery program in high elevation lakes of California’s Sierra Nevada. Over 45,000 fish were removed by means of gillnets and electrofishing, with length-at-capture and date-at-capture recorded for nearly every captured fish ^11,12^. After every fishing event, we estimated the selection differential on fish length as the difference in the population mean trait value before and after selection. This differential is equal to the covariance between the trait and relative fitness ^9,10^, which can be partitioned into the product of three components: the correlation between the trait (body length) and fitness (captured = 0, not-captured = 1), the variability (standard deviation) of fitness, and the variability (standard deviation) of the trait. We then related among-population variation in these components of selection to among-population variation in lake physical characteristics (lake surface, maximum depth, elevation), demographic (fish length, population size, density), fishing gear (gillnet, electrofishing), and fishing intensity (proportion of the population captured, number of fishing events, and time until extirpation) (see Extended Table 1).

### Repeatable yet variable fishery selection

In most populations, the largest fish were quickly and consistently removed by fishing, which dramatically altered the population size structure. Both mean fish length and its variability decreased with increasing cumulative catch (Figure 1). On average, fish length decreased by −1.03 mm ± 0.13 mm (SD) for each additional percent of the population removed (i.e., “mm/%”; linear mixed regression: t_33.2_ = −7.86, p < 0.001). The average shift in mean length from the start to the end of the fish removal period was ~100 mm (45% of initial mean body length). Fish length variability (here measured as SD – similar results were obtained for CV) also decreased with increasing cumulative catch: 0.32 mm ± 0.07 mm/% (linear mixed regression: t_25.8_ = −4.60, p < 0.001). These estimates are conservative because the mean duration of fish removal efforts was 2.5 years, during which time survivors would continue to grow. With a moderate growth correction (5% per year, see Methods), the estimates were −1.11 ± 0.13 mm/% for mean length (linear mixed regression: t_33.4_ = −8.68, p < 0.001) and −0.35 ± 0.06 mm/% for variability in length (linear mixed regression: t_25.3_ = −5.4, p < 0.001). With a larger – but still plausible – growth correction (10% per year), the estimates were 1.18 ± 0.13 mm/% for mean length (linear mixed regression: t_33.2_ = −9.37, p < 0.001) and −0.37 ± 0.06 mm/% for variability in length (linear mixed regression: t_25.1_ = −6.06, p < 0.001). These decreases could not be explained by random harvesting because permutation simulations, in which mortality was random, i.e., independent of fish length, yielded non-significant slope coefficients for mean length (Figure 2a, one-sample t-test: t_99_ = −1.173, p = 0.25) and much lower slope coefficients for length variability (Figure 2b, one-sample t-test: t_99_ = −9.54, p < 0.01).

**Figure 1:**
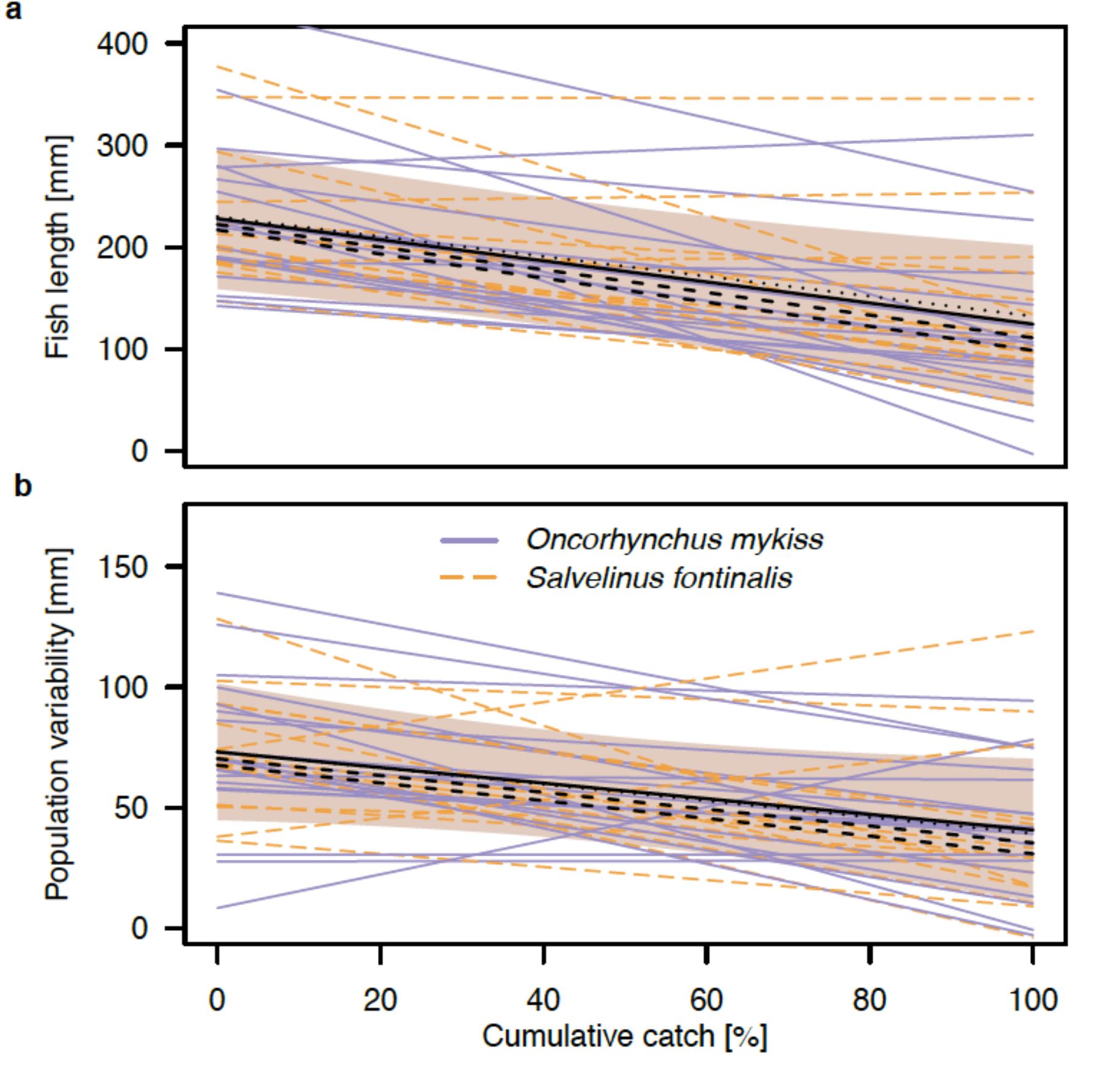
Effect of size-selective fishing on population structure. _Plots demonstrate length-at-capture in mm (p < 0.001)(a) and surviving population’s length variability (standard deviation) (p < 0.001) (b) as a function of the percentage of the population removed to date (i.e., the ‘catch intensity’). Thin lines represent the response of individual populations (linear regressions), and bold lines represent the average response of both *Salvelinus fontinalis* (brook trout; dashed orange) and *Oncorhynchus mykiss* (rainbow trout and hybrids; solid purple) populations (mixed-model). Black dashed lines represent the average response using 5% and 10% growth corrections, and black dotted lines represent the response with correction for reproduction after the initiation of the experimental fisheries (see Methods). Shaded area represents mean estimates ± standard deviation. Analyses were based on 43,827 individuals in 35 populations in (a), and 1092 standard deviation estimates in(b) (population details reported in Extended Table 1).

**Figure 2:**
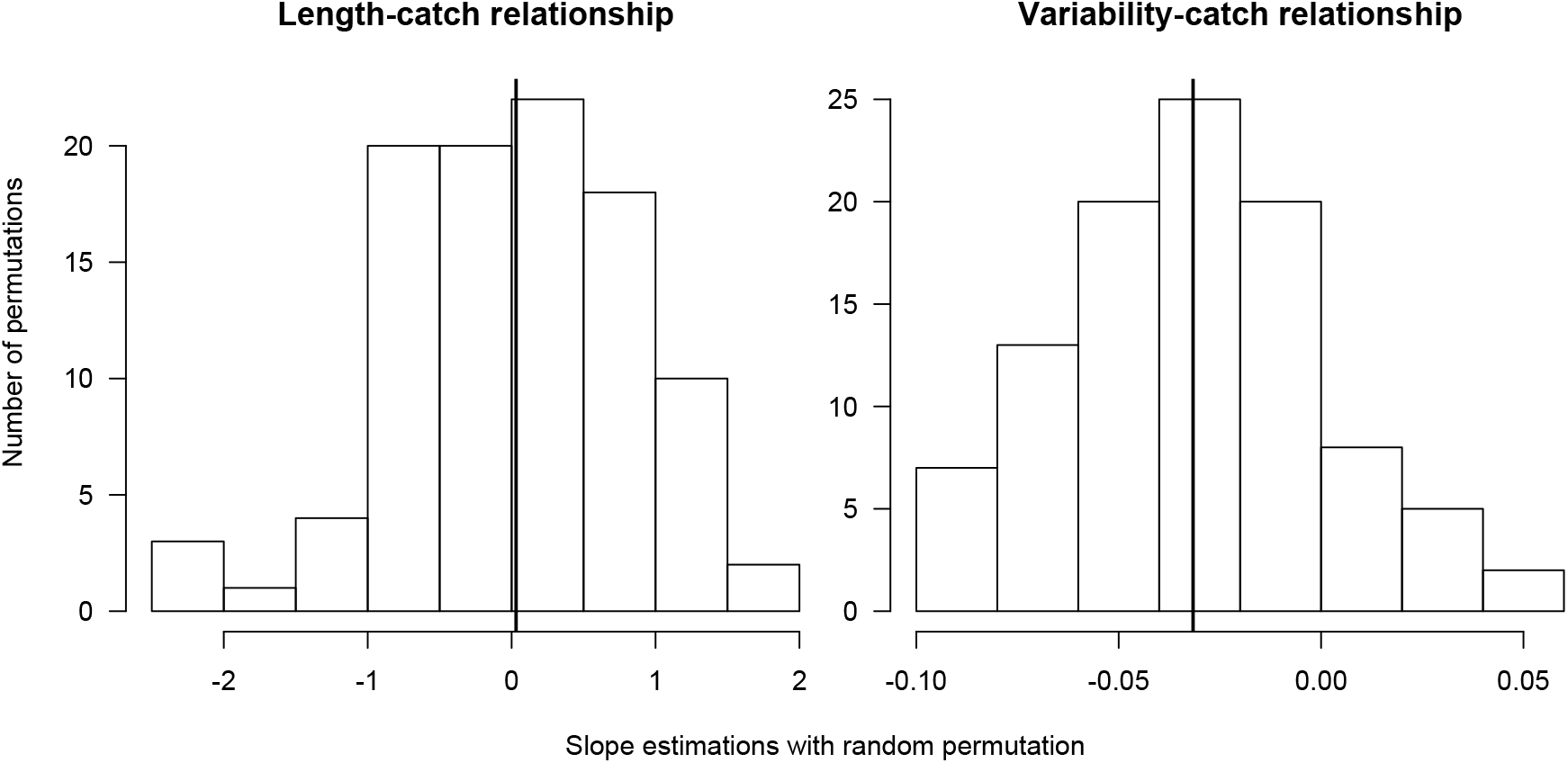
Empirical distributions of the slopes of the length-catch and variability-catch relationships inferred from random permutations of the date at-capture. These distributions represent the expected values of the relationship if fishing is random with respect to size (100 random permutations). Random fishing is not expected to alter the size structure of the population (left-hand panel, slope of the length-catch relationship is 0.08 ± 1.00, not significantly different from zero). Random fishing is expected to slightly reduce variability (right-hand panel, slope of variability-catch relationship = −0.03 ± 0.04), but less than non-random fishing where the slope of the variability-catch relationship is expected to be ten times larger (0.32 mm ± 0.07 mm/%).

The negative association between mean length and cumulative catch yielded increasingly negative selection differentials (i.e., acting against large fish) with increasing cumulative catch (Figure 3). Most differentials were negative (918 of 1092), with the few positive estimates tending to occur when precision was low (i.e., when few individuals remained in the population; Figure 3). The magnitude of the selection differential increased with increasing cumulative catch: −0.31 ± 0.14 mm/% (mixed-model linear regression: t_30.2_ = −2.20, p < 0.05). As above, these effects were even stronger when length was corrected for growth during the study period: −0.35 ± 0.13 mm/% with a 5% correction (linear mixed regression: t_30.2_ = − 2.6, p < 0.05); and −0.39 ± 0.12 mm/% with a 10% correction (linear mixed regression: t_30.4_ = −3.17, p < 0.01). Importantly, even moderate fishing intensity (30–40% of the population captured) led to selection differentials greater than 10 mm. That is, fish surviving to that level of fishing were, on average, 10–20 mm (6–13%) smaller than those in the original population (Figure 3). These estimates are similar to other freshwater fish populations subject to low or moderate fishing intensity, where selection differential estimates on individual growth tend to be 5–17% ^13,14^.

**Figure 3:**
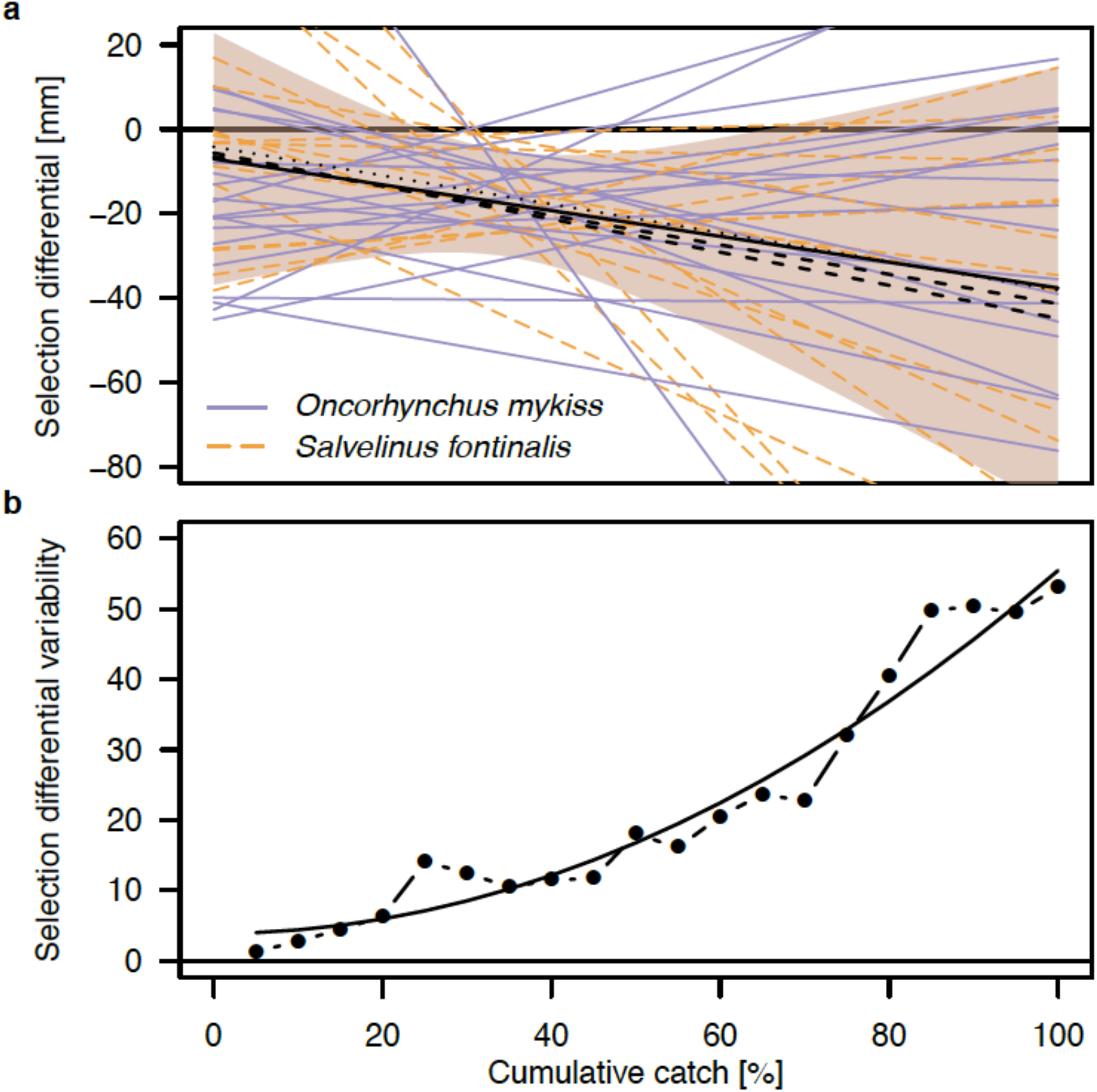
Effect of cumulative catch on the selection differential. Plots demonstrate the selection differential, that is, the difference in mean length between the initial population (i.e., before fishing) and the survivors (i.e., after fishing) (linear mixed model, p < 0.05) (a) and selection differential variability (standard deviation) (quadratic model, p < 0.001) (b) as a function of the percentage of the population removed to date (i.e., catch intensity). Thin lines represent the responses of individual populations, and bold lines represent the average response of both *Oncorhynchus mykiss* (rainbow trout and hybrids; solid purple) and *Salvelinus fontinalis* (brook trout; dashed orange) populations. Dashed black lines represent the average response using 5% and 10% growth corrections, and black dotted lines represent the response with correction for reproduction after the initiation of the experimental fisheries (see Methods). We used 1092 selection differentials estimates from 35 populations in (a), and one standard deviation estimate per 5% fraction of catch in (b) (population details reported in Extended Table 1).

Among-population variability in selection differentials increased dramatically at harvest levels above 80% (Figure 3). Indeed, modeling the distribution of selection differentials after binning fishing intensity into 5% intervals revealed that the variability of selection differentials (standard deviation within each 5% interval) increased non-linearly with fishing pressure (quadratic, R^2^ = 0.62, F_1,17_ = 30.3, p < 0.001). These results suggest that the outcome of size-selective fishing becomes increasingly unpredictable when catch levels are high (>80%), with some populations exhibiting large decreases in mean length and other populations not responding or even exhibiting increases in mean length.

These results confirm that fishing-induced selection can modify population size structure, even at low fishing pressures. More importantly, these results also highlight how the consequences of selection can be highly variable among replicate populations. This unpredictability could explain some of the contradictory results of fishing-induced selection reported in the literature ^4^. Moreover, our result that size-selective fishing becomes increasingly unpredictable with higher fishing intensity provides further evidence that reducing total harvest might be the best action to recover severely impacted populations ^15^.

### Multiple factors mediate fishery selection

The outcomes of fishing-induced selection differed among populations, which provided an opportunity to explore the relative importance of the three components of selection and examine how each was affected by environmental or demographic attributes, or the fishing regime.

One component of selection is the correlation between the trait and fitness, in our case represented by the selectivity of the fishing gear. Our results suggest that this correlation is the most important component of selection in our system, accounting for 60.9% of the variance in selection differentials (linear regression: F_1,33_ = 51.5, p < 0.001, Figure 4). Variation among populations in this correlation, and hence selection differentials, could be explained by several factors. First, correlations between length and fitness were stronger in populations with initially smaller fish (proportion of variance (PoVE) in the correlation explained by mean fish length: 29.6%, F_1,33_ = 13.9, p < 0.001). Second, correlations were stronger in populations with higher densities (PoVE = 14.7%, F_1,33_ = 5.68, p < 0.05). Third, correlations were weaker when the lake had at least one tributary (PoVE = 11.6%, F_1,33_ = 4.34, p < 0.05). These associations indicate that in a closed environment with many relatively small fish, relatively large fish are more likely to be caught earlier. This suggests that the selectivity of fishing gear (i.e., the strongest selective force) is highly influenced by the environment. Indeed, a multiple regression including all three parameters (fish length, density, and tributary presence) explained 47.6% of the variance in correlation between trait and fitness (Figure 4).

**Figure 4:**
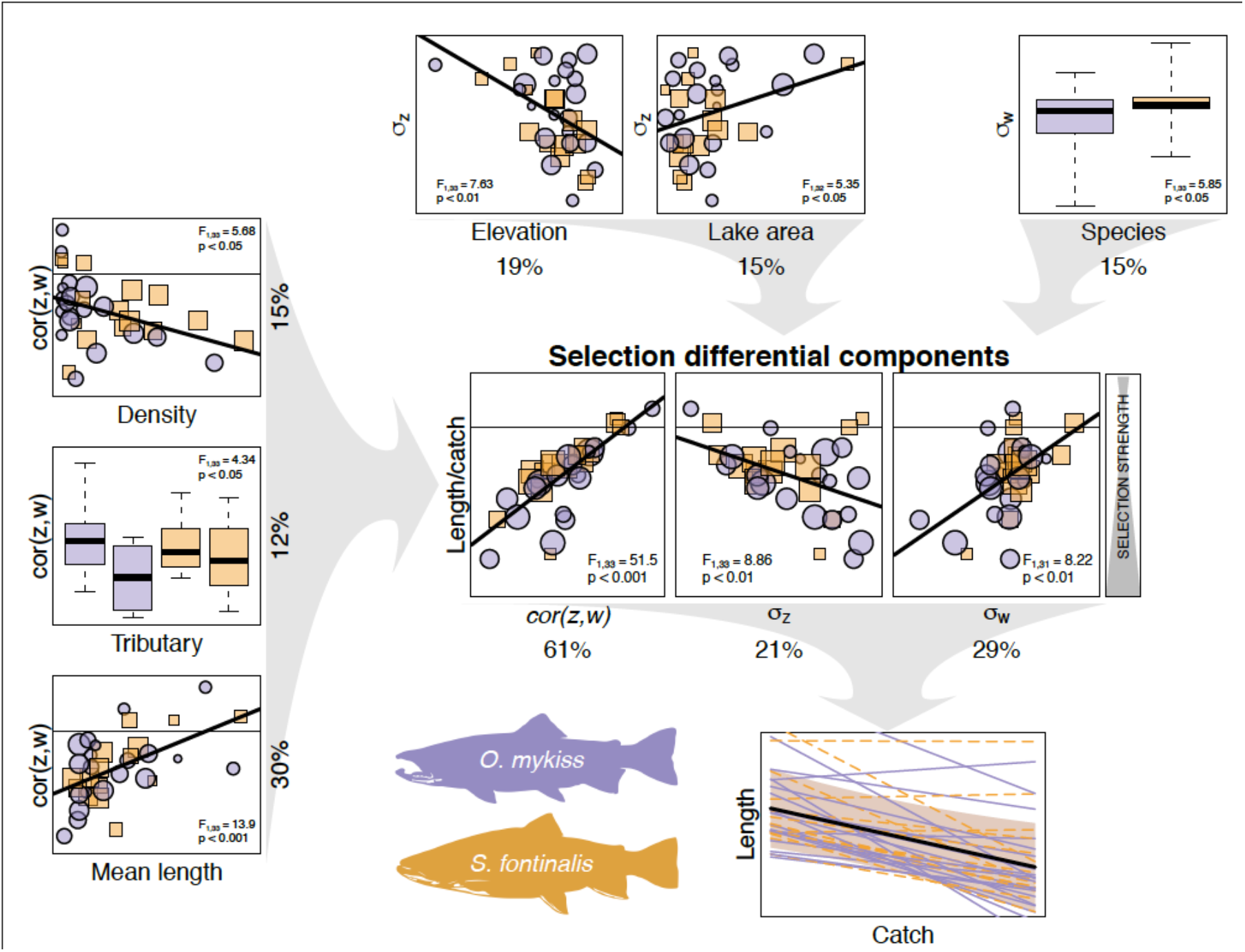
Schematic to show the main drivers of fishing-induced selection. The length-catch relationship (bottom-right) represents selection, as expressed in Figure 1a. The three central plots represent the selection differential components (fitness-trait correlation, trait variability, fitness variability) and the relationship between each component and the length-catch relationship, orange circle for rainbow trout and purple square for brook trout (symbol size is proportional to log-transformed population size). The regression statistics are included on each plot. The percentage displayed below each plot represents the univariate proportion of variance explained (note that these do not add up to 100%, as they can be correlated). Using a multivariate model, the proportion of variance would be: 44% for the correlation, 21% for trait variability, and 17% for fitness variability. The six remaining plots (on the left and top) represent the influence of environmental, demographic, and species-specific factors on the selection components. The proportion of variance in a multivariate model explaining the correlation between trait and fitness (left-hand plots) is 29.6% for mean length, 13.8% for the presence of a tributary, and 4.2% for density, and the proportion of variance in a multivariate model explaining the length variability (top-left) is 18.8% for elevation, and 8.6% for lake area.

Another component of selection is variability of the trait (i.e., size), which here accounted for 21.1% of the variance in selection among populations (F_1,33_ = 8.86, p < 0.01, Figure 4). Here too the environment played a role; length was more variable in populations at lower elevations (PoVE = 18.8%, F_1,33_ = 7.63, p < 0.01) and in larger lakes (PoVE = 15.2%, F_1,33_ = 5.93, p < 0.05). These results imply that, in our system, lakes with greater habitat suitability (i.e., lower elevations are associated with warmer temperatures and a longer ice-free season ^16^) and complexity (i.e., larger systems tend to exhibit more complex habitat structure ^17^) support a more diverse size structure, which makes them more susceptible to selection. This result might provide insight into conflicting results in the literature, where some ^13,14^ – but not all 18 – studies report strong selection against large fish in heavily-fished populations. That is, even strong gear selectivity can yield only weak selection if trait variability is already low – a hypothesis that would be easy to test in ongoing fisheries.

The last component of selection is variability in fitness, which here accounted for 29.3% of the among-population variance in selection (F_1,33_ = 13.7, p < 0.001, Figure 4), with populations experiencing evenly-distributed fishing events throughout the study being less susceptible to fishery selection. In our study system, fitness variability is indeed directly linked to the fishing regime, with populations experiencing constant (even) fishing mortality throughout the season exhibiting lower fitness variability than populations subjected to sporadic (uneven) fishing events (see Extended Figure 1). This final component was not obviously linked to any environmental factors; yet it was the only factor that differed between species, with rainbow trout exhibiting slightly lower fitness variability than brook trout (PoVE = 15.1%, F_1,33_ = 5.85, p < 0.05), potentially due to species-specific behavior around fishing gears ^19,20^.

### Management Implications

Decomposing selection into its three components provides an opportunity to identify the most important component of selection and, simultaneously, the appropriate target for management. Our results suggest that overall selection can be influenced differentially by each of the three selection components, each associated with different management strategies to reduce the unintended consequences of fishing-induced selection. In our particular system, even with gear that is substantially less selective than those used in modern-day fisheries ^21,22^, the most important factor influencing the strength of fishing-induced selection was the correlation between fitness and length, i.e., the fishing gear selectivity. In order to manage populations against the potentially negative effect of size-selective fishing, managers could introduce fishing gear regulations such as population-specific mesh combinations protecting the largest individuals ^23,24^. In addition, our data suggest that tributaries entering lakes may provide some protection against size-selective fishing. Tributaries may provide refugee for larger fish (i.e., a “reduced take zone”) that decreases susceptibility to size-selective fishing. This supports the argument that reserves provide an important management tool to mitigate fishing-induced selection for populations in which selection is mostly driven by fishing gear selectivity ^25–28^.

Alternatively, in populations where selection is mostly driven by trait variability, management measures could focus on increasing the variability in fish length through the protection of larger fish. Our results further suggest that environmental characteristics such as habitat suitability and diversity also influence trait variability, suggesting that habitat restoration activities could be implemented as an indirect path to promoting trait variability ^29^.

Finally, where selection is driven by fitness variability, management measures that target fishing schedules and intensity (e.g., duration of the fishing season, daily yield) could be effective strategies for reducing fishing-induced selection. Our results suggest that fitness variability is directly linked to the fishing schedule (see Extended Figure 1), with even fishing throughout the season being less selective than infrequent, but intense, fishing events ^30^.

These findings also highlight an interesting feedback that might shape fisheries-induced selection, and our ability to detect it. In particular, strong size-selectivity of fishing gear imposes substantial selection on body size in harvested fishes. At the same time, however, this selectivity should decrease size (and fitness) variability, which should reduce the intensity of selection. Therefore, we have a situation where selection differentials might decrease through time in harvested populations even if gear selectivity itself is constant. Stated more generally, selection differentials might not be good indicators of gear selectivity, and investigating how the different components of selection vary through time might be necessary to assess how size-selective fishing has impacted populations.

### Conclusions

Our study helps explain why the outcomes of fishing-induced selection reported in the literature are contradictory ^4^. First, size-selective fishing becomes increasingly unpredictable with higher fishing intensity. Second, responses to fishery selection are population-specific because fishery selection is influenced by demographic- and habitat-specific factors. Predicting population responses to fishing-induced selection is a key challenge faced by fishery managers ^31–33^, but it has been overshadowed by controversy surrounding its effects and underlying mechanisms ^34,35^. By working with replicated populations, we were able to explore practical principles of fishing-induced selection that could help guide management of harvested stocks. Most importantly, evolutionary impact assessments ^31^ should replicate selection measures at the subpopulation level to investigate the repeatability of selection across time and space. Doing so will enable managers to disentangle the components of selection and to tailor their management actions accordingly.

Overall, our study demonstrates that fishing is highly selective, even at low harvest rates. As size-related traits are heritable ^36^, this selection has the potential to dramatically modify both the size and genetic structures of harvested populations. However, the outcomes of selection are highly variable and can be modified by environmental, demographic- and species-specific factors. Disentangling these factors and the different components of selection (fitness-trait correlation, trait variability, fitness variability) is critical to producing flexible and relevant management strategies, but does rely on repeated measurements of selection. However, such experiments may not need vast numbers of individuals because our results suggest that selection differentials varied linearly with catch. Evolutionary impact assessments that incorporate these components of selection will allow managers to predict how a population will respond to different harvest rates and management scenarios with far greater accuracy than is currently achieved.

## Methods

### Study system: large-scale experimental fisheries

The Sierra Nevada mountain range contains thousands of lakes that are naturally fishless above 1800 meters ^37^. Most of these lakes have been stocked with non-native trout, some as early as 1850, to create recreational fishing opportunities ^38^. By 1996, 63% of the lakes larger than one hectare were inhabited by non-native populations of rainbow trout (including hybrids with golden trout) and/or brook trout ^38^. In the past 20 years, research has revealed detrimental effects of non-native fish on native biota, particularly endemic mountain yellow-legged frogs (*Rana muscosa* and *R. sierrae*) ^11,16,39^. Consequently, several large-scale fish removal programs were initiated in Inyo National Forest and in Sequoia, Kings Canyon, and Yosemite National Parks. These programs involved sinking multimesh (six panels, each with a different mesh size from 10–38 mm) gillnets (86% of all captures) and performing electrofishing on lake shores and inlet and outlet streams (14%) ^11^. The multimesh nets are designed to target fish across the full range of sizes and ages, and are generally considered less selective than traditional fishing gear ^23,40^.

To date, fish have been completely removed from over 60 locations (lakes and rivers); and from these, we analyzed lake populations where at least 15 fish were captured over at least three fishing events. This reduced dataset included 35 populations from 32 lakes: 17 lakes with rainbow trout, 12 lakes with brook trout, and three lakes with both species. The number of fish removed per lake ranged from 19 to 4741 over 29 to 3307 days (Extended Table 1). A total of 47,679 fish were caught and the total length-at-capture (mm) and date-at-capture were recorded. The dataset includes almost all the fish present in each lake, excluding only fish that died from natural mortality during the removal period.

The 32 lakes exhibited large variability in environmental and population-specific parameters (Extended Table 1). Lake size was indexed as lake perimeter (207–2118 m), lake area (1989–124625 m^2^), and maximum depth (2.5–30 m). Lake quality (i.e., the potential for rapid fish growth) was indexed as elevation (2146–3583 m; lower elevations assumed to represent higher quality habitat ^16^) and maximum fish length (200–550mm). Population size structure was indexed as mean fish length (range among lakes: 93–347 mm) and length variability (standard deviation; range among lakes: 27–94 mm). Lakes also differed in population size (19–4741 individuals), density (5–3355 individuals per ha), and various aspect of the fishing regime, including the proportion of fish caught with gillnets (28–100%) versus electrofishing, the number of fishing events (5–114 events), and the mean number of fish caught per event (2–145 fish per event) or per day during the overall study duration (0.05–6.9 fish per day).

### Catch and length

We assessed the selective effect of fishing by correlating fish length with catch intensity. Specifically, after every fishing event, we assigned to each fish a catch intensity value equal to the number of fish caught to date, divided by the total number of fish in the lake (i.e., the proportion of fish captured to date). We then assessed the relationship between catch intensity and length-at-capture by means of linear mixed-models; these models included length-at-capture as response variable, catch intensity and species as a fixed effects, and lake ID as a random effect, allowing the slope and intercept to vary among lakes. We used the same type of model to assess the effect of fishing intensity on the variability of length-at-capture, which was measured after each fishing event, as the standard deviation of the length of all the fish remaining in the lake. Analyses were performed using both linear and logarithmic relationships, as changes over time might not be linear. The linear model had the better fit and so was used in all analyses presented herein. Because no differences emerged between the two species (rainbow trout and brook trout), we pooled species for all analyses presented here (Figure 1); we tested for species effects when analyzing the component of selection (see below and Figure 4).

For the sake of simplicity, we first assumed that fish did not grow during the study period (typically 2–3 years), which would result in a conservative (underestimated) estimate of selection. This simplification is reasonable given that fish growth is extremely low in these high elevation lakes due to low temperatures and productivity, as well as the short growing season ^16^. However, to examine the sensitivity of our results to growth, we next adjusted the observed length-at-capture at a given capture date by expected growth rate since the start of the experimental fisheries. To do so, we applied two growth scenarios: a slow growth correction assuming 5% length increase per year and a fast growth correction assuming 10% length increase per year. With this method, the length of the fish captured during the course of removal was adjusted to their expected length at the first date of sampling after allowing for growth.

To test if random sampling could explain the observed selective effects of fishing - estimated as the slopes of the length-catch intensity relationships (Figure 1a), and variability-catch intensity relationships (Figure 1b) - we performed 100 simulations where mortality occurred randomly; i.e., we used random permutations of the time-at-capture of each fish. We performed the same tests (linear mixed models) on the random simulations, allowing direct comparison of the observed and simulated data. For each simulation, we extracted the slope of the relationship between length-at-capture (or length variability) and catch intensity, in order to build an empirical distribution of the expected change in length (and variability) with catch. We assumed that these distributions were normally distributed, estimated their means and standard deviations, and tested if the expected change (based on the mean of the distributions) differed from zero with one-sample t-tests (Figure 2).

When possible, fish were prevented from reaching spawning areas in the associated streams by makeshift dams and/or gillnets to block access to inlet and outlet streams. In addition, efforts were made to reduce reproduction during the experimental fisheries, by searching for and destroying redds (groups of fertilized eggs buried in spawning grounds). However, some reproduction cannot be completely ruled out after the initiation of the experimental fisheries, in particular for brook trout that are known to breed in lake habitat ^41^. If any reproduction occurred, it likely occurred during the first year following removal initiation and likely produced few fish as most adults were removed quickly. To estimate the potential effect of reproduction after the initiation of the fisheries, we performed a sensitivity analysis by repeating the analysis after removing fish smaller than 50 mm that were caught after the first year of survey, i.e., 334 fish potentially born after removal initiation. This is a conservative correction, as we also removed all the smaller fish born before the start of the study. Re-analyzing the data using this reduced dataset, the results were nearly identical, with a corrected relationship between length and catch of −97.7 ± 13.1 [***], i.e., a 5% difference, which is less than the difference with moderate growth correction (see above). The corrected relationship between variability and catch was −0.34 ± 0.06 [***], and the corrected relationship between selection differential and catch was −0.34 ± 0.15 [*], see Fig. 1 & 3 for the reported impact. Thus, fishing-induced selection appears to be driven by the removal of the largest fish, and the effects of potential reproduction during the program were deemed to be negligible.

### Estimating selection differentials

For each lake, we compared the mean length of surviving fish after each fishing event (1092 events) to the mean length before the same fishing event, yielding a selection differential for each event in each lake. Under several common assumptions, this differential is equal to the covariance between trait and relative fitness ^10^. Using this covariance, we can disentangle the different components of selection into the products of (1) the correlation between the trait and fitness, (2) the variability (standard deviation) of fitness, and (3) the variability of the trait under selection (see below) ^10^.

Selection differential: 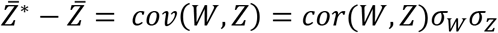

With 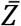 the average value of the trait Z in the population (in this study Z is the length of the fish), 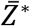 the average value of the trait Z after selection, i.e., in the sub-population that survived the harvest event, W the relative fitness, *σ_W_*, the standard deviation of the relative fitness, and *σ_Z_* the standard deviation of the length in the initial population.

We evaluated fishing-induced selection by correlating selection differentials with catch intensity. Specifically, for each lake we compared the selection differential generated by each fishing event to the proportion of fish captured to date (i.e., the catch intensity, described above). We then assessed the relationship between selection differentials and catch intensity by means of linear mixed-models; these models included selection differentials as the response variable, catch intensity as a fixed effect, and lake ID as a random effect, allowing the slope and intercept to vary among lakes (Figure 3a).

To assess if catch intensity also affected the variability in selection differentials, we binned catch intensity into 5% intervals and estimated, for each lake, the average selection differential in each 5% interval. For example, to estimate the average selection differential in the 15–20% interval, we averaged - for each lake - all the selection differential estimations with catch intensity being within this interval. Then we measured the between lake variability within each interval as the standard deviation of the averaged selection differentials. Since there is only one value per catch interval, we used linear models to assess how the variability in selection differentials was affected by catch intensity (Figure 3b).

### Selection differential components and environmental parameters

In each lake and after each fishing event, we estimated the three components of selection. Specifically, after each fishing event, fish that had been captured were assigned a fitness value of zero, and one if not captured. This allowed estimating (1) the correlation between body length and fitness, the standard deviation of fitness (*σ_W_*), and the standard deviation in body length before selection (*σ_Z_*) in each lake and after every fishing event. To allow for between-lake comparisons and to estimate the effect of attributes related to the environment, fish population, or the fishing regime, we calculated, for each selection component, a single estimate per lake. For the covariance and the correlation between fitness and trait, we used the mean value of all estimates (calculated after every fishing event). To estimate the standard deviation in fitness in each lake (*σ_W_*), we used the standard deviation of all individuals’ relative time-at-capture (as a proportion of the population captured). Specifically, within each lake, we transformed the time-at-capture into a [0,1] variable, with zero being assigned to the fish caught during the first fishing event, and one assigned to the fish caught during the last fishing event. This value represents how spread the fishing events are throughout the study duration, and is minimal when several fishing events are distributed evenly throughout the study and maximal when few fishing events are distributed at the beginning and the end of the study (see Extended Figure 1). We then estimated the standard deviation in body length before selection (*σ_Z_*). Finally, because the selection differential is the covariance between fitness and trait, and is mathematically linked to the three components of selection components ^10^, we calculated an independent value for the selection differential. Instead of the covariance between fitness and trait, we used the relationship between catch and length-at-capture (see Figure 1a) as a proxy for overall selection. This value represents the magnitude of length change when catch increases and can be understood as fishing-induced selection. We assessed the validity of this proxy with a linear regression between the slope of the length-catch relationship and the covariance between length and fitness (linear regression: F_1,33_ = 14.3, p < 0.001, R^2^ = 0.30).

Finally, we assessed the contribution of each selection component to overall selection by means of weighted analysis of variance. Specifically, overall selection was the response variable, the three components of selection were predictor variables (Figure 4), and the logarithm of population size was the weighting parameter. We used one-way analysis of variance for each individual component, but also checked the additive effects of selection components with multiple regression. Finally, we assessed the contribution of the environmental and population-specific variables (lake size, lake quality, population structure, population size, and fishing regime) to each selection component by means of stepwise linear regression and weighted analysis of variance as before.

### Data and statistics

The data that support the findings of this study belong to the National Park Service and the California Department of Fish and Wildlife, and can be accessed through them upon reasonable request. The code used to generate results of this study is available from the corresponding author upon request.

All analyses were performed in R ^42^. The exact sample size in each group is reported in Extended Table 1, for both the number of individuals per lake (Pop. size) and for the number of selection estimates generated after each fishing event (Number of fishing events). Effects are reported as mean ± standard deviation for linear models, and percentage of variance for analyses of variance. All test are two-sided, and reported with the value of the statistic and the degrees of freedom. Model assumptions of residual normality were assessed visually with histograms. Mixed models were fitted with the lmer0 function from the package lme4 ^43^. Significance tests were computed according to Kuznetsova et al. ^44^ using the package lmerTest.

## Acknowledgments

We thank the dozens of individuals who have been involved in the removal of invasive trout in the Sierra Nevada as well as the National Park Service (particularly Ninette Daniele, Travis Espinoza, and Daniel Boiano) and California Department of Fish and Wildlife (particularly Mitch Lockhart and Sarah Mussulman) for sharing the data collected. SN was funded by the Swiss National Science Foundation (P2LAP3_148434) and the Parrotia Foundation and APH was supported by a Visiting Miller Professorship from the Miller Institute at UC-Berkeley.

## Authors contributions

SN and SMC conceived the project. SN analyzed the data. SN, APH, AMS, and SMC discussed the analyses and wrote the manuscript. RAK and MTB collected data and provided feedback on the manuscript.

## Extended Figure 1 – Catch distribution

**Extended Figure 1:**
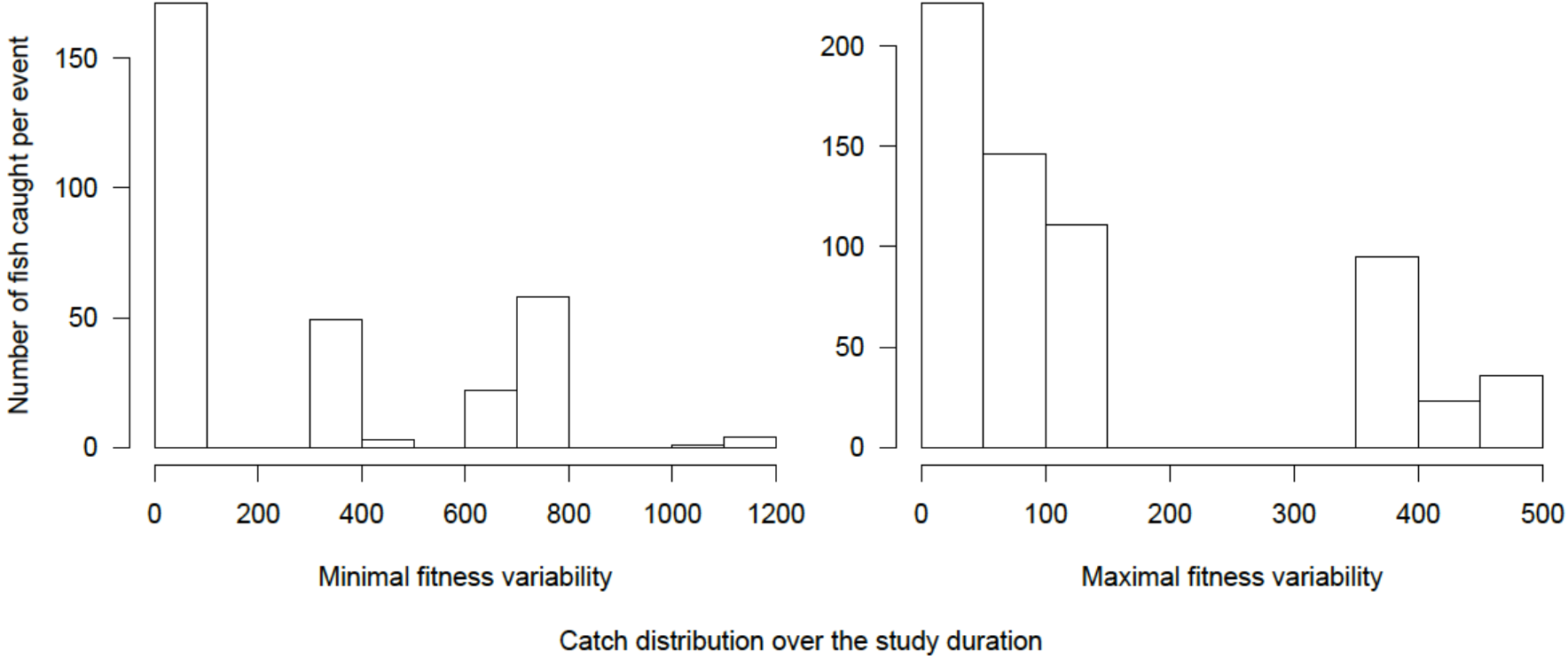
representations of the two most extreme catch distributions determining the fitness variability. For the population on the left-hand panel, the fitness variability was minimal and fishing was relatively evenly distributed throughout the study period. For the population on the right-hand panel, the fitness variability was maximal, and fishing events were distributed at the start and end of the study.

## Ext. Table:Environmental and population specific parameters

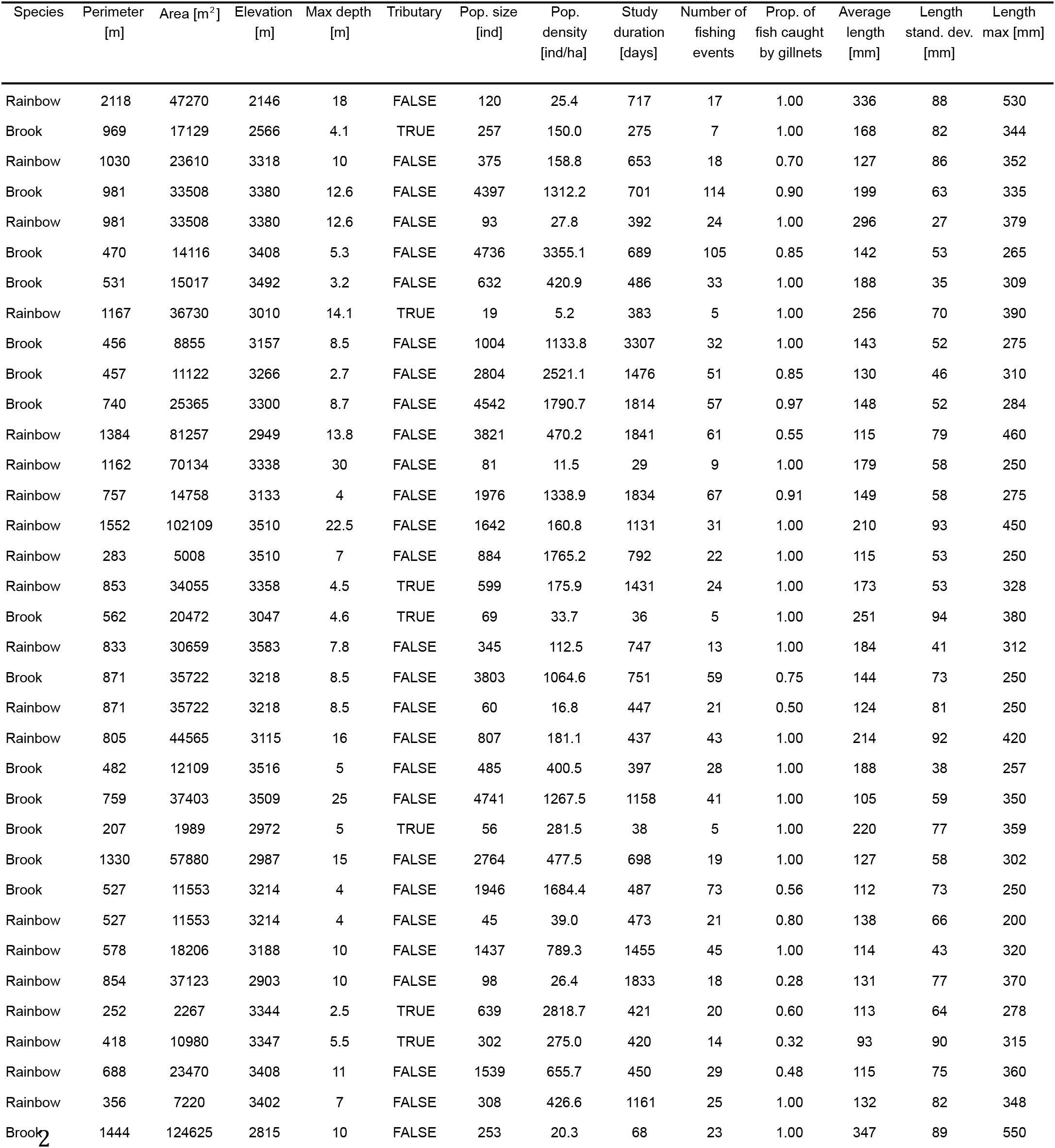

